# RAB1B interacts with TRAF3 to promote antiviral innate immunity

**DOI:** 10.1101/542050

**Authors:** Dia C. Beachboard, Moonhee Park, Madhuvanthi Vijayan, Dillon J. Fernando, Graham D. Williams, Stacy M. Horner

## Abstract

Nucleic acid-based antiviral innate immunity activates a signaling cascade that induces type I and type III interferons (IFNs), and other cytokines. This signaling, which is highly regulated, is initiated by pattern recognition receptors, such as RIG-I, that sense viral RNA and then signal to the adaptor protein, MAVS. This adaptor protein then recruits additional signaling proteins, including TRAF3 and TBK1, to form a signaling complex that results in IRF3 activation for transcriptional induction of IFN. Here, we show that the GTPase trafficking protein RAB1B positively regulates RIG-I signaling to promote IFN-β induction and the antiviral response. Over-expression of RAB1B increases RIG-I-mediated signaling to IFN-β, while deletion results in reduced signaling of this pathway. Additionally, this loss of RAB1B results in a dampened antiviral response, as Zika virus infection is enhanced in the absence of RAB1B. Importantly, we identified the mechanism of RAB1B action by determining that it interacts with TRAF3 to facilitate the interaction of TRAF3 with MAVS. Thus, we identified RAB1B as a regulator of TRAF3 to promote the formation of innate immune signaling complexes in response to nucleic acid sensing.

---

Viruses are detected in the infected cell by the antiviral innate immune system. This system is activated when specific pattern recognition receptors sense pathogen-associated molecular patterns (PAMPs) that are unique to viruses for self-nonself discrimination. These viral PAMPs include cytosolic nucleic acids derived from either RNA or DNA viruses. Specifically, RIG-I and MDA5 sense viral RNA in the cytoplasm, while cGAS and IFI16 sense viral DNA (1,2). Upon sensing of viral nucleic acids, these sensor proteins become activated, allowing them to signal to their respective adaptor proteins, MAVS and STING (3,4). These adaptors then recruit the downstream signaling molecules that ultimately drive the transcriptional induction of type I and type III interferons (IFNs) (5), leading to the production of hundreds of IFN-stimulated genes (ISGs), many of which have antiviral functions (6).

The antiviral innate immune response is carefully regulated to adequately inhibit viral infection without causing excessive inflammation in host tissues. This regulation can occur in several ways, including post-translational modification of signaling proteins, unique protein-protein interactions between signaling proteins and their regulators, and through changes in localization of signaling proteins and regulators to different subcellular compartments. The RNA sensor RIG-I is regulated by all three of these mechanisms. Before viral infection, RIG-I resides in the cytoplasm in an inactive state (7). However, during RNA virus infection, both Riplet and TRIM25 ubiquitinate RIG-I with K63-linked ubiquitin chains (8–10). This ubiquitination allows RIG-I to interact with the chaperone protein 14-3-3ε. Subsequently, 14-3-3ε translocates RIG-I into membranes to interact with MAVS and induce downstream signaling (11). Besides RIG-I, other antiviral signaling proteins are regulated by localization changes during induction of antiviral innate immune signaling. For example, the serine/threonine kinase TBK1 and the E3 ubiquitin ligase TRAF3 both have been shown to relocalize upon nucleic acid sensing (12,13) to interact with the MAVS signaling complex at mitochondria and ER contact sites, otherwise known as mitochondrial-associated ER membranes (MAMs) (14). The mechanism(s) by which these proteins relocalize upon nucleic acid sensing and the consequences of this relocalization are not fully understood.

Previously, we have found that during viral infection many cellular proteins change their localization to and from MAMs (15). One of these proteins is the GTPase protein RAB1B, which is recruited into MAMs and the MAVS signaling complex during RIG-I pathway activation (15). RAB1B, which is functionally distinct from the related RAB1A protein, regulates ER-to-Golgi trafficking by binding to effector proteins that tether ER-derived vesicles to the cytoskeleton to facilitate the movement of these vesicles to the Golgi (16,17). Interestingly, a known RAB1B effector protein called p115 has previously been shown to interact with TRAF3 and is required for TRAF3 trafficking from the Golgi to the MAVS signaling complex (13), suggesting that RAB1B may be required for TRAF3 interaction with MAVS during RIG-I signaling.

Here, we have identified a role for RAB1B in positively regulating RIG-I pathway signaling to type I IFN. We show that RAB1B interacts with TRAF3 to promote the interaction of TRAF3 with the signaling adaptor protein MAVS. Ultimately, this work reveals that the known cellular trafficking protein RAB1B interacts with TRAF3 to facilitate the assembly of innate immune signaling complexes, providing a new example of a trafficking protein that is repurposed to regulate the host response to virus infection.

## RESULTS

### RAB1B positively regulates RIG-I pathway signaling to IFN-β and the antiviral response

Previously, we found that following RIG-I signaling RAB1B localizes to the MAM and is in a complex with MAVS (15). To determine if this MAVS-interacting protein also regulates RIG-I/MAVS pathway signaling to IFN-β, we measured Sendai virus (SenV)-mediated signaling to the IFN-β promoter following over-expression of RAB1B in 293T cells. SenV is a negative-strand RNA virus that potently induces RIG-I/MAVS signaling to IFN-β (18,19). We found that while over-expression of RAB1B on its own did not significantly induce signaling to IFN-β, it did augment SenV-mediated signaling to IFN-β, as measured by an IFN-β promoter luciferase assay, and it increased *IFNB1* mRNA levels, as measured by real time quantitative PCR (RT-qPCR) (Fig. 1A-B). Importantly, over-expression of RAB1B also enhanced the levels of *IFNB1* transcripts following transfection of a RIG-I agonist (the hepatitis C virus (HCV) 5’ppp-containing polyU/UC RNA) (20), suggesting that RAB1B directly regulates RIG-I pathway signaling (Fig. 1C). We also found that RAB1B over-expression enhanced SenV-mediated induction of *IFNB1* transcripts in the human monocyte THP1 cells (Fig. 1D). Collectively, these data suggest that while RAB1B does not induce signaling on its own, it does enhance RIG-I pathway signaling to IFN-β.

**Figure 1.**
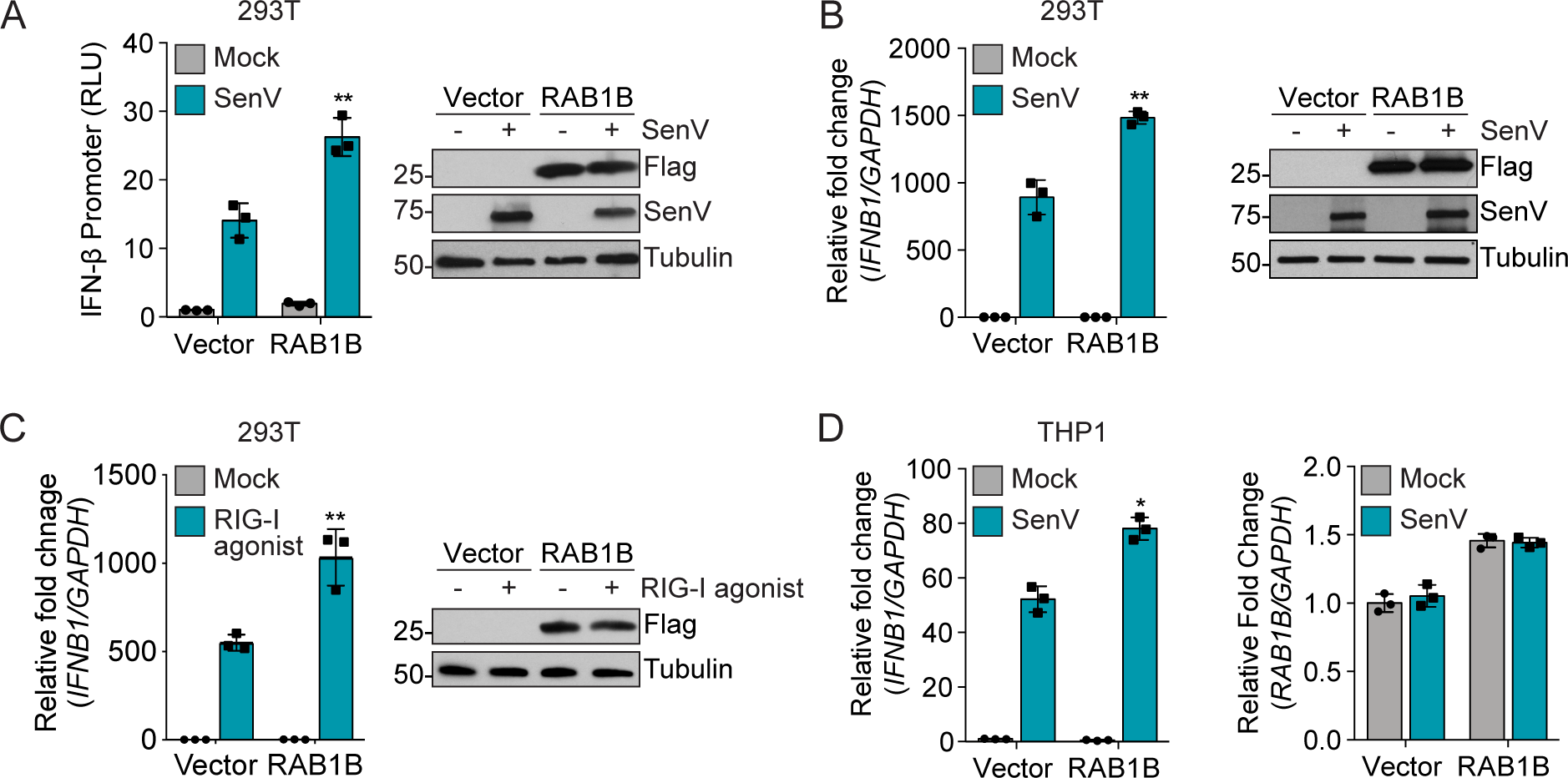
RAB1B positively regulates RIG-I pathway signaling to IFN-β. *A*, IFN-β promoter reporter luciferase expression from 293T cells transfected with indicated plasmids and then Mock-or SenV-infected (16 h). RLU, relative luciferase units. Immunoblot for Flag-RAB1B, SenV, and Tubulin. *B-D,* RT-qPCR analysis of RNA from 293T (*B-C*) or THP1 (*D*) cells expressing Flag-RAB1B or Vector and stimulated with SenV (12 h) or a RIG-I agonist (8 h). *IFNB1* transcript levels were measured relative to *GAPDH* and normalized to the Vector, Mock condition. RAB1B expression was verified by immunoblot analysis of extracts from paired samples in (*A-C*) and by RT-qPCR for *RAB1B* relative to *GAPDH* in (*D*). Individual dots represent technical replicates with bars displaying the mean ± SD (n=3) of one of three representative experiments. * P≤0.05, ** P≤0.01, by an unpaired Student’s t-test comparing the activated samples.

To determine whether loss of RAB1B decreased RIG-I pathway signaling to IFN-β, we depleted RAB1B in 293T cells using an siRNA. Depletion of RAB1B resulted in a decrease in the induction of *IFNB1* transcripts following SenV infection (Fig. 2A). To confirm this result, we generated 293T cells deficient in RAB1B by using CRISPR/Cas9 and then measured SenV-induced signaling to IFN-β by RT-qPCR in the parental or RAB1B knockout (KO) 293T cells. Loss of RAB1B resulted in decreased induction of *IFNB1* transcripts following SenV infection, confirming that RAB1B positively regulates IFN-β induction (Fig. 2B).

**Figure 2.**
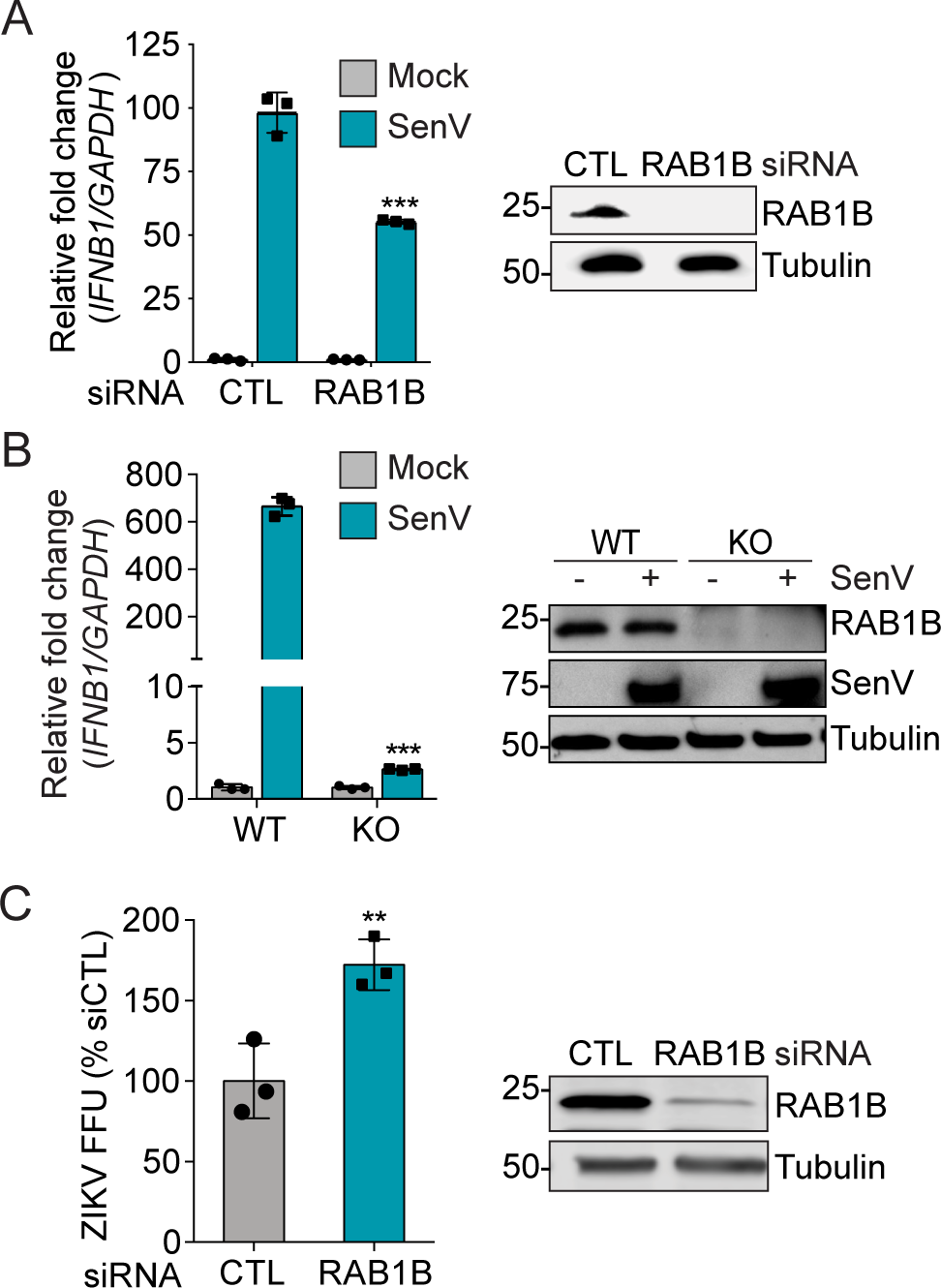
RAB1B is required for full induction of IFN-β in response to RIG-I pathway activation by SenV. *A*, RT-qPCR analysis of *IFNB1*, relative to *GAPDH* and normalized to the Vector, Mock sample, in 293T cells treated with control (CTL) or RAB1B siRNAs for 24 h and then Mock-or SenV-infected (8 h), along with immunoblot analysis for RAB1B expression. *B*, RT-qPCR analysis for *IFNB1*, relative to *GAPDH* and normalized to the WT, Mock sample, in WT or RAB1B KO 293T cells infected with SenV (20 h). For *A* and *B*, individual dots represent technical replicates with bars displaying the mean ± SD of one of three representative experiments. *C,* Focus forming assay of ZIKV focus-forming units (FFU) from supernatants of Huh7 cells infected with ZIKV (48 hours post infection, MOI 0.01) after transfection of CTL or RAB1B siRNAs (48 h), measured as the percentage of FFU relative to siCTL. Individual dots represent biological replicates with bars displaying the mean ± SEM (n=3). ** P≤0.01, *** P≤0.001, by an unpaired Student’s t-test comparing SenV samples (*A-B*) and or siRAB1B to siCTL (*C*).

Since RAB1B positively regulated induction of IFN-β, we hypothesized that it would also be required for antiviral responses driven by IFN. Therefore, we tested if loss of RAB1B resulted in higher levels of viral infection during Zika virus (ZIKV) infection, a virus known to be susceptible to type I IFN (21). We treated Huh7 cells with either an siRNA to RAB1B or control, infected these cells with ZIKV (multiplicity of infection (MOI) 0.01), and then 48 hours later measured the production of infectious ZIKV particles in the supernatant using a focus-forming assay. In RAB1B depleted cells, ZIKV titer was increased by approximately fifty percent as compared to the control cells (Fig. 2C). Collectively, these data reveal that RAB1B positively regulates the antiviral response.

### RAB1B is required for phosphorylation of IRF3 and TBK1

The induction of IFN-β in response to viral infection requires the kinase TBK1, which is activated by autophosphorylation. This activated TBK1 then phosphorylates the transcription factor IRF3 at multiple residues, allowing its translocation to the nucleus where it cooperates with NF-κB to transcriptionally induce IFN-β (5,22,23). To determine if RAB1B was required to activate this signaling cascade, we first measured the phosphorylation of IRF3 at S396 (p-IRF3) in response to SenV infection in the parental and RAB1B KO 293T cells by immunoblot analysis. We found that the phosphorylation of IRF3 in response to SenV was decreased in the RAB1B KO cells as compared to the parental cells, revealing that RAB1B activates innate immune signaling upstream of IRF3 phosphorylation (Fig. 3A). Next, to test whether RAB1B is required for the activation and autophosphorylation of TBK1 in this system, we measured phosphorylation of TBK1 at S172 (p-TBK1) after SenV infection in the parental and RAB1B KO 293Ts by immunoblot analysis. We found that TBK1 phosphorylation in response to SenV was decreased in the RAB1B KO cells as compared to the parental cells (Fig. 3B). Taken together, these data suggest that RAB1B promotes signaling that leads to TBK1 autophosphorylation and subsequent IRF3 phosphorylation in response to RIG-I activation.

**Figure 3.**
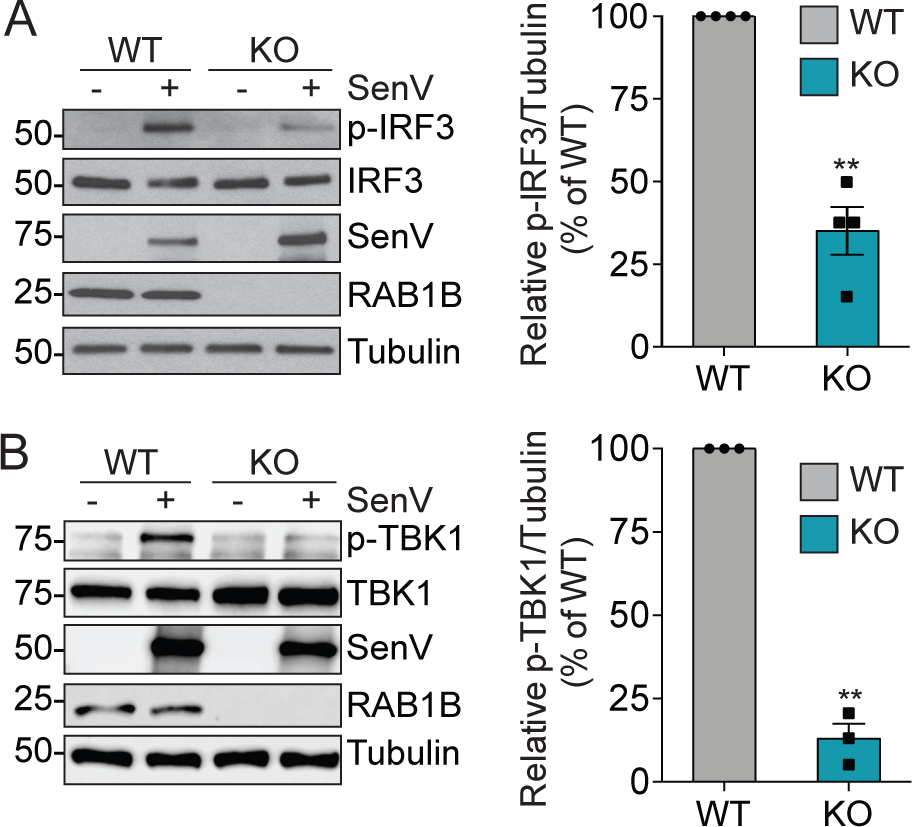
RAB1B is required for phosphorylation of IRF3 and TBK1 in response to RIG-I pathway activation by SenV. *A-B*, Immunoblot analysis of WT and RAB1B KO 293T cells that were Mock-or SenV-infected (*A*, 20 h; *B*, 6 h). Right panels show quantification of p-IRF3 or p-TBK1 induction, relative to Tubulin, in response to SenV, with the WT condition set to 100%. Individual dots represent biological replicates (n = 3-4), with the bars displaying the mean ± SEM. ** P≤0.01, by an unpaired Student’s t-test.

### RAB1B interacts with TRAF3

As RAB1B is required for TBK1 activation, this suggested to us that RAB1B may act on a protein that regulates TBK1. The TRAF proteins, specifically TRAF2, TRAF3, and TRAF6, are well-known regulators of TBK1 activation (24). We decided to focused on TRAF3 specifically because the known RAB1B effector protein p115 has previously been shown to regulate TRAF3 function during RIG-I signaling (13). Therefore, to test if RAB1B interacted with TRAF3, we over-expressed HA-RAB1B and Myc-TRAF3 in cells in which we activated RIG-I signaling by over-expression of the constitutively active form of RIG-I (RIG-I-N) (25). We found that Myc-TRAF3 co-immunoprecipitated with HA-RAB1B only upon activation of the RIG-I pathway (Fig. 4A). We have previously shown that RAB1B is in complex with MAVS following MAVS overexpression, which is known to activate signaling (15). Therefore, to determine if MAVS signaling is sufficient for RAB1B to interact with TRAF3, we over-expressed HA-RAB1B and GFP-TRAF3, as well as Flag-MAVS to activate signaling, in cells. We found that GFP-TRAF3 co-immunoprecipitated with HA-RAB1B only in the presence of over-expressed MAVS, which was also detected in the complex, suggesting these proteins form a complex in response to signaling (Fig. 4B).

**Figure 4.**
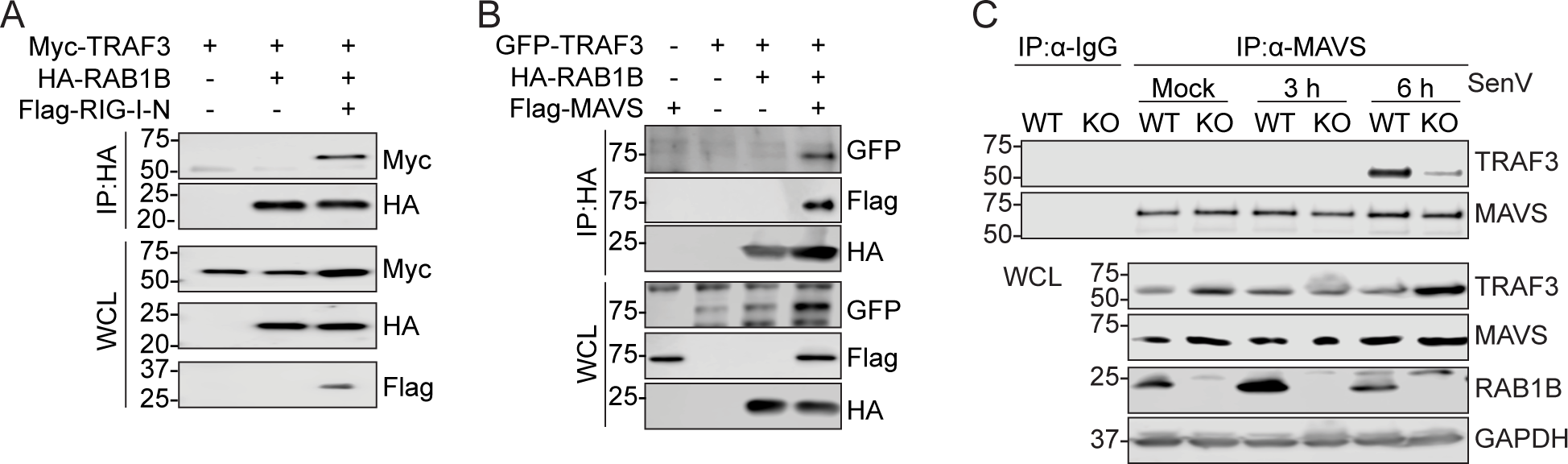
RAB1B interacts with TRAF3 during RIG-I signaling. *A*, Immunoblot analysis of anti-HA immunoprecipitated extracts and whole cell lysates (WCL) from 293T cells expressing HA-RAB1B, Myc-TRAF3, and Flag-RIG-I-N. Representative of three biological replicates. *B*, Immunoblot analysis of anti-HA immunoprecipitated extracts and WCL from 293T cells expressing HA-RAB1B, GFP-TRAF3, and Flag-MAVS. Representative of three biological replicates. *C*, Immunoblot analysis of anti-MAVS immunoprecipitated (or anti-IgG) extracts and WCL from WT or RAB1B KO 293T cells that were Mock-or SenV-infected for the indicated times. Representative of three independent experiments.

### RAB1B is required for TRAF3 to interact with MAVS

As TRAF3 interaction with MAVS is required for TBK1 activation and recruitment to the MAVS signaling complex (26,27), we next tested if RAB1B was required for this TRAF3-MAVS interaction. We measured the interaction of endogenous MAVS and TRAF3 during a time course of SenV-infection by co-immunoprecipitation. We found that MAVS and TRAF3 interacted at six hours post-SenV infection in 293T cells; however, this interaction was reduced in the RAB1B KO 293T cells (Fig. 4C). These data demonstrate that RAB1B facilitates the interaction of TRAF3 with MAVS in response to activation of innate immune signaling.

## DISCUSSION

Although the signaling proteins in cytosolic nucleic acid sensing pathways have been well studied, the molecular mechanisms by which these proteins are regulated are less defined. Previously, we identified proteins that may regulate the assembly of the MAVS signaling complex at membrane contact sites between ER, mitochondria, and peroxisomes through proteomic analysis of the proteins that relocalize into these contacts sites during RIG-I activation (15). Some of these proteins with differential membrane association upon RIG-I signaling included GTPase proteins, such as RAB1B. Here, we have shown that RAB1B positively regulates RNA sensing by promoting RIG-I signaling to IFN-β through interactions with the E3 ubiquitin ligase TRAF3. This interaction facilitates TRAF3 recruitment to MAVS, leading to phosphorylation of TBK1 and IRF3 for the transcriptional induction of IFN-β. In summary, our work reveals that the known cellular trafficking protein RAB1B interacts with TRAF3 to facilitate the assembly of the MAVS signaling complex, providing a new example of a trafficking or chaperone protein that is repurposed to regulate the host response to virus infection (see model in Fig. 5).

**Figure 5.**
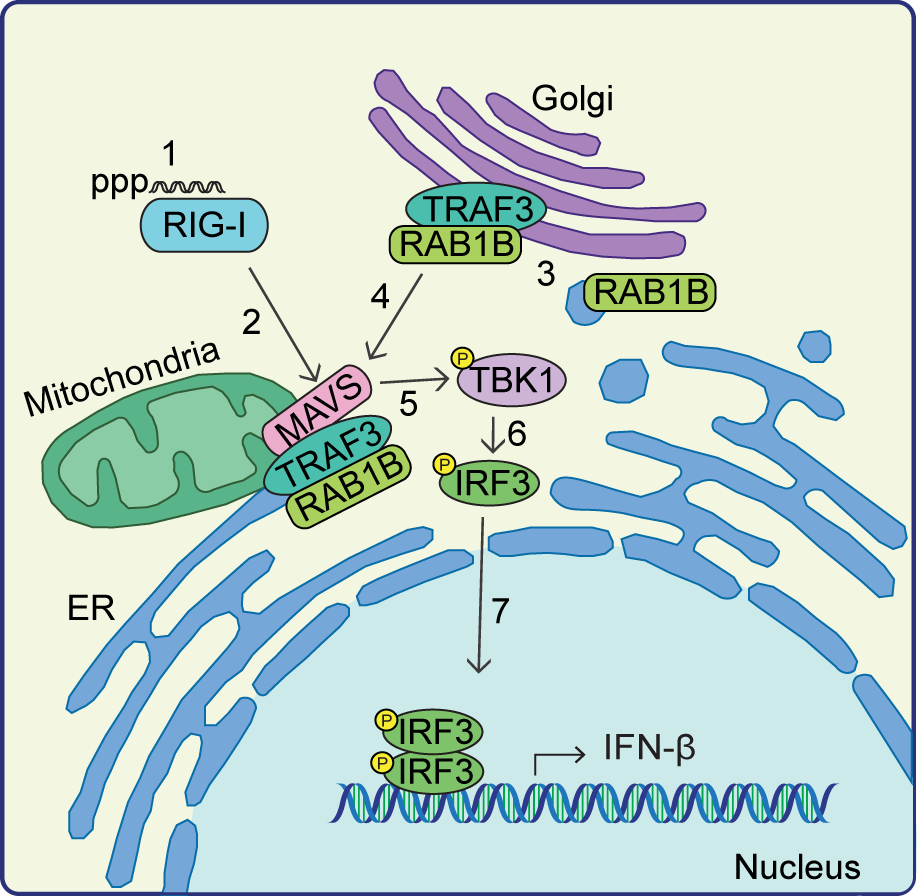
Model of RAB1B regulation of IFN-β-induction. RNA viruses are sensed in the cytoplasm by RIG-I or MDA5 (Step 1). After sensing viral RNA, RIG-I translocates to ER-mitochondrial contact sites at mitochondrial-associated ER membranes where it interacts with MAVS (Step 2). RAB1B, normally involved in ER-Golgi trafficking, then interacts with TRAF3 (Step 3) and facilitates TRAF3 interaction with MAVS (Step 4). This interaction may be facilitated by re-arrangements in intracellular membranes of the Golgi and/or ER. TRAF3-MAVS interaction results in activation and autophosphorylation of TBK1 (Step 5), which then phosphorylates IRF3 (Step 6), resulting in IRF3 dimerization and translocation to the nucleus, where it cooperates with NF-κB to induce IFN-β transcription and the subsequent antiviral response (Step 7).

It is unclear if RAB1B directly regulates TRAF3 or if RAB1B effector proteins are required for this regulation. We know that small GTPase proteins such as RAB1B regulate intracellular membrane trafficking by interacting with effector proteins (16). RAB1B, similar to other GTPase proteins, cycles between an inactive GDP-bound state and an active GTP-bound state that associates with intracellular membranes (16). Importantly, this GTP-bound, membrane-associated RAB1B facilitates COPI recruitment and cargo transport between the ER and the Golgi by interactions with specific effector proteins (28). These effector proteins include tethering factors that link organelles and vesicles prior to fusion, adaptors for motor proteins that direct organelle trafficking, and regulators of other GTPases that are recruited to the specific subcellular compartments of those GTPases (29). Therefore, RAB1B effector proteins are critical for RAB1B trafficking functions and are likely important for mediating the role of RAB1B in antiviral innate immunity. Indeed, the RAB1B effector protein p115 has previously been shown to interact with TRAF3 and be required for TRAF3 recruitment to the MAVS signaling complex (13). Thus, it is likely that RAB1B interacts with p115 to facilitate the movement of TRAF3 on vesicles from the Golgi apparatus to site of MAVS signaling. There is evidence that Golgi and other intracellular membranes rearrange during MAVS signaling to bring MAVS in contact with the Golgi-associated TRAF3 (13,14). As we were unable to detect relocalization of RAB1B to MAVS-signaling sites (data not shown), this suggests that only a small proportion of RAB1B, likely the p115-bound form, interacts with TRAF3 at any given time. Indeed, RAB1B has multiple effector proteins (16,30), which suggests that different pools of RAB1B in the cell may have different functions that could be activated by interactions with specific effector proteins.

The mechanisms by which RAB1B would interact with a unique set of effectors during cytosolic nucleic acid sensing are not entirely clear. While the overall localization of RAB1B does not change in the cell during innate immune signaling, it is possible that specific post-translational modifications are altered (either added or removed) on RAB1B, which may facilitate interactions with a new set of effectors. Indeed, post-translation modification of small GTPase proteins does alter their association with effector proteins and therefore their biological functions. For example, the GTPase RALB is ubiquitinated by K63-ubiquitin linkages to switch from its role in autophagy to a role in innate immune signaling (31,32). This ubiquitination prevents RALB from binding to its autophagy effector protein EXO84 and instead allows RALB to bind to the Sec5 effector protein for interaction with TBK1 to regulate innate immunity. Similarly, ubiquitination of the ARF domain of TRIM23 activates the GTP hydrolysis activity of TRIM23 to regulate the function of TBK1 in autophagy (33). Therefore, it is possible that changes in post-translational modification of RAB1B allows it to interact with TRAF3 for innate immune signaling. Other innate immune signaling proteins are also known to be activated by specific post-translational modifications (34). Therefore, addition and removal of post-translational modifications on signaling chaperone or trafficking proteins may be a general cellular mechanism to repurpose these proteins into innate immune signaling regulators, and future studies will examine this possibility for RAB1B.

Our results suggest that the interaction of RAB1B with TRAF3 is important to induce RIG-I/MAVS-mediated innate immune responses. We do not yet know if RAB1B interacts only with TRAF3 or if it also regulates the functions of TRAF2 and TRAF6. As TRAF2, TRAF3, and TRAF6 have been shown to act at different time points after signaling induction (27), it is possible that while RAB1B regulates TRAF3, other RAB or GTPase proteins may regulate TRAF2 or TRAF6 to promote innate immune signaling.

Our work adds RAB1B to a growing list of GTPase proteins that have been shown to regulate innate immune signaling, either by regulating trafficking of innate immune signaling proteins or specific effector protein interactions. For example, cytosolic DNA-induced innate immune responses mediated by cGAS have been shown to be regulated by RAB2B and its effector protein GARIL5 (35). Signaling from the bacterial LPS sensor TLR4 is also regulated by RAB proteins (36–39). Specifically, both RAB10 and RAB11A positively regulate TLR4 signaling (36), with RAB10 promoting TLR4 recycling to the plasma membrane, and RAB11B promoting TLR4 localization to the *Escherichia coli* phagosome (38,39). Conversely, RAB7B negatively regulates TLR4 signaling by promoting lysosomal degradation of TLR4 (37). As intracellular innate immunity is so often regulated at the cell biological level (40), it is likely that other RAB proteins will control specific intracellular trafficking events that regulate nucleic acid-induced innate immunity and viral infection.

## EXPERIMENTAL PROCEDURES

### Cell culture

293T, Huh7, and Vero cells were grown in Dulbecco’s modification of Eagle’s medium (DMEM; Mediatech) supplemented with 10% fetal bovine serum (FBS; HyClone), and 25 mM 4-(2-hydroxyethyl)-1-piperazineethanesulfonic acid (HEPES; Thermo-Fisher). THP1 cells (gift from Dr. Dennis Ko, Duke University, who obtained them from the American Type Culture Collection (ATCC)) were grown in Roswell Park Memorial Institute medium 1600 (RPMI; Thermo-Fisher) supplemented with 10% FBS (HyClone), and 25 mM HEPES (Thermo-Fisher). The identity of the Huh7 cells in this study were verified using the GenePrint STR Kit (Promega, Duke University DNA Analysis Facility). The 293T and Vero (CCL-3216 and CCL-81) cells were obtained from ATCC; the Huh7 cells were a gift of Dr. Michael Gale at the University of Washington. Cells were verified as mycoplasma-free using the LookOut PCR detection Kit (Sigma).

### Virusess

SenV Cantell strain was obtained from Charles River Laboratories and used at 200 hemagglutination units/mL. SenV infections were performed in serum-free media for 1 h, after which complete media was replenished. ZIKV-Dakar (DAK; Zika virus/A.africanus-tc/SEN/1984/41525-DAK) (GenBank accession #KU955591) was provided by Dr. Scott Weaver at University of Texas Medical Branch. Stocks were prepared as described (41). ZIKV infections were performed at a MOI of 0.01 for 48 h in Huh7 cells.

### Focus forming assay for ZIKV titer

Supernatants were harvested from ZIKV-infected cells at 48 hours post infection, serially diluted, and used to infect naïve Vero cells in triplicate wells of a 48-well plate for 2 h before overlay with methylcellulose. After 48 h, plates were fixed in methanol acetone. Cells were blocked (10% FBS in PBS) and then immunostained with the mouse anti-4G2 antibody, generated from the D1-4G2-4-15 hybridoma cell line against the flavivirus Envelope protein (ATCC). Infected cells were visualized following incubation with a horseradish peroxidase-conjugated secondary antibody (1:500; Jackson ImmunoResearch) and VIP Peroxidase Substrate Kit (Vector Laboratories). Titer (focus-forming units (FFU)/mL) was calculated from the average number of 4G2 positive foci at 10X magnification, relative to the amount and dilution of virus used.

### Plasmids

The following plasmids have been previously described: pEF-TAK-Flag (42), pEF-TAK-Flag-RAB1B (15), pEF-BOS-Flag-RIG-I-N (25), pIFN-β-luc (43), pGL4.74 [hRluc/TK] (Promega), pcDNA-Blast (44), and plEGFP-N1-TRAF3 (gift from Dr. Soman Abraham, Duke University) (45). The following plasmids were generated during this study: pEF-TAK-HA-RAB1B, pcDNA-Myc-TRAF3, pX330-sgRAB1B. pEF-TAK-HA-RAB1B was generated by PCR amplification of RAB1B from pEF-TAK-Flag-RAB1B and insertion into the *EcoRI-XbaI* digested pEF-vector using InFusion cloning (Clontech). The pcDNA-Myc-TRAF3 plasmid was generated by cloning the PCR-amplified TRAF3 (plEGFP-N1-TRAF3) into pcDNA-Myc using *SalI* and *KpnI*. To generate the CRISPR guide RNA plasmid, sgRNA oligonucleotides were annealed and inserted into the *BbsI*-digested pX330 (46). The oligonucleotide sequences used for cloning are listed in Table 1. The plasmid sequences were verified by DNA sequencing and are available upon request.

**Table 1:**
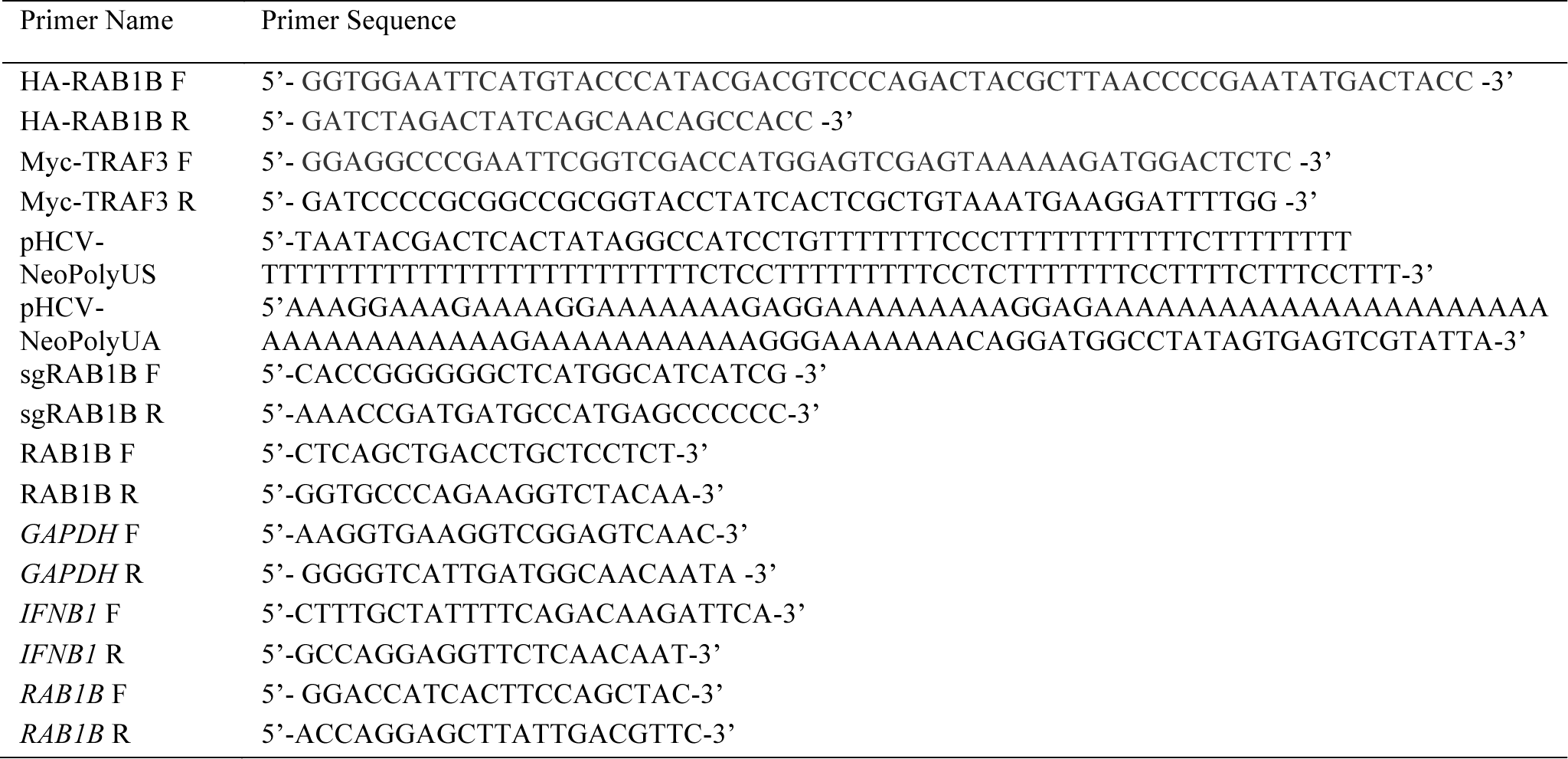
Primers used for cloning and RT-qPCR.

### Generation of RIG-I agonist

Annealed oligonucleotides containing the sequence of the HCV 5’ppp poly-U/UC region (Table 1) (20) were in vitro transcribed using the Megashortscript T7 transcription kit (Ambion) followed by ethanol precipitation, with the resulting RNA resuspended at 1 µg/µL for use in experiments.

### Transfection

DNA transfections were performed using FuGENE6 (Promega). RIG-I agonist transfections were done using the TransIT-mRNA Transfection kit (Mirus Bio). The siRNA transfections were done using Lipofectamine RNAiMax (Invitrogen). The IFN-β-promoter luciferase assays were performed as previously described at 12 or 16 h post Mock or SenV infection and normalized to the Renilla luciferase transfection control (18).

### Generation of KO cells lines

RAB1B KO 293T cells were generated by CRISPR/Cas9, using a single guide targeting exon 4 designed using the CRISPR design tool (http://crispr.mit.edu). pX330-sgRAB1B was transfected into 293T cells along with pcDNA containing blasticidin resistance (pcDNA-Blast) for 24 h. Cells were then replated at limiting dilutions into 15 cm dishes with 10 µg/mL blasticidin treatment for 3 days. Single cell colonies were then amplified and screened for RAB1B expression by immunoblot. Genomic DNA was extracted from candidate RAB1B KO cells using QuickExtract DNA Extraction Solution (Lucigen), as per manufacturer’s instructions. This genomic DNA was then amplified (using primers across exon 4, see Table 1), cloned into the pCR4-TOPO TA vector (Invitrogen), and Sanger sequenced. Eleven resulting genomic DNA subclones were sequenced, and nine clones had a 27 bp deletion at the end of exon 4 that extended into the intron and two clones had a 1 bp insertion in exon 4 that resulted in a frame shift that lead to a premature stop codon within exon 5.

### RNA Analysis

Total cellular RNA was extracted using the Purelink RNA mini kit (Life Technologies). RNA was then reverse transcribed using the iScript cDNA synthesis kit (BioRad), as per the manufacturer’s instructions. The resulting cDNA was diluted 1:3 in ddH_2_O. RT-qPCR was performed in triplicate using the Power SYBR Green PCR master mix (Thermo-Fisher) and the Applied Biosystems Step One Plus or QuantStudio 6 Flex RT-PCR systems. The oligonucleotide sequences used are listed in Table 1.

### Immunoblotting

Cells were lysed in a modified radioimmunoprecipitation assay (RIPA) buffer (10 mM Tris [pH 7.5], 150 mM NaCl, 0.5% sodium deoxycholate, and 1% Triton X-100) supplemented with protease inhibitor cocktail (Sigma) and Halt Phosphatase Inhibitor (Thermo-Fisher), and post-nuclear lysates were isolated by centrifugation. Quantified protein (between 5-15 µg) was resolved by SDS/PAGE, transferred to nitrocellulose or polyvinylidene difluoride (PVDF) membranes in a 25 mM Tris-192 mM glycine-0.01% SDS buffer and blocked in StartingBlock (Thermo-Fisher) buffer, 5% milk in phosphate-buffered saline containing 0.01% Tween-20 (PBS-T), or 3% BSA in Tris-buffered saline containing 0.01% Tween-20 (TBS-T). After washing with PBS-T or TBS-T (for phosphoproteins) buffer, membranes were incubated with species-specific horseradish peroxidase-conjugated antibodies (Jackson ImmunoResearch, 1:5000) followed by treatment of the membrane with ECL+ (GE Healthcare) or Clarity Western ECL substrate (BioRad) and imaging on X-ray film or a LICOR Odyssey FC. The following antibodies were used for immunoblotting: R-anti-RAB1B (Santa Cruz, 1:1000), R-anti-SenV (MBL, 1:1000), M-anti-Flag M2 (Sigma, 1:5000), R-anti-GFP (Thermo-Fisher, 1:1000), M-anti-Tubulin (Sigma, 1:5000), R-anti-p-IRF3 (Cell Signaling Technology, 1:1000), M-anti-IRF3 ((47), 1:1000), R-anti-p-TBK1 (Cell Signaling Technology, 1:1000), R-anti-TBK1(Cell Signaling Technology, 1:1000), anti-HA (mouse-Abcam and rabbit-Sigma, 1:1000), anti-TRAF3 (mouse-Santa Cruz or rabbit-Cell Signaling Technology, 1:1000), anti-MAVS (mouse-AdipoGen or rabbit-Bethyl Laboratories, 1:1000), R-anti-GAPDH (Cell Signaling Technology, 1:1000), and anti-Myc (mouse-Santa Cruz or rabbit-Cell Signaling Technology, 1:1000).

### Quantification of immunoblots

Immunoblots imaged using the LICOR Odyssey FC were quantified by ImageStudio software, and raw values were normalized to relevant controls for each antibody. Immunoblots developed on film, were quantified using FIJI, which normalizes the signal to relevant controls for each antibody (48). ImageStudio and FIJI give similar quantification results when compared directly. Phosphoprotein values were normalized to Tubulin and displayed as the percentage of signal from WT.

### Immunoprecipitation

Cells were lysed as above but in a modified RIPA buffer for immunoprecipitation (25 mM Tris [pH 7.5], 150 mM NaCl, 1% sodium deoxycholate, 10% glycerol, and 1% Triton X-100). Quantified protein (between 200-500 µg) was incubated with protein-specific or isotype control antibody (Cell Signaling Technology) in lysis buffer at 4°C overnight with head over tail rotation. The lysate/antibody mixture was then incubated with Protein G Dynabeads (Invitrogen) for 2 h. Beads were washed 3X in PBS or modified RIPA buffer for immunoprecipitation and eluted in 2X Laemmli Buffer (BioRad) supplemented with 5% 2-Mercaptoethanol at 95°C for 10 min. Proteins were resolved by SDS/PAGE and immunoblotting, as above.

### Statistical Analysis

Student’s unpaired t-test was used for statistical analysis of the data using GraphPad Prism software. Graphed values are presented as mean ± SD or SEM (n = 3 or as indicated); *p ≤ 0.05, **p ≤ 0.01, and ***p ≤ 0.001.

## Acknowledgements

We thank all members of the Horner lab for discussion and reading of the manuscript. We thank the following people for reagents and equipment use: Dr. Michael Gale Jr. of University of Washington, Dr. Scott Weaver of University of Texas Medical Branch, Dr. Dennis Ko of Duke University, Dr. Soman Abraham of Duke University, and the Duke Functional Genomics Core Facility. This work was supported by funds from the NIH: K22 AI100935 (S.M.H.); T32-CA009111 (D.C.B.). Additional funding sources include the American Cancer Society Postdoctoral fellowship 131321-PF-17-188-01-MPC (D.C.B.); the Burroughs Welcome Fund (S.M.H); and a Duke Bridge Award (S.M.H).

